# Social context modulates cannibalistic attraction in *Drosophila* larvae

**DOI:** 10.64898/2026.01.08.698439

**Authors:** Amelie Edmaier, David Walter, Nora Tutas, Roxana Probst, Katrin Vogt

**Affiliations:** Department of Biology, University of Konstanz, Konstanz, Germany; Max Planck Institute of Animal Behavior, Konstanz, Germany

## Abstract

Animals often face conflicting decisions to ensure survival. Under food limitation, cannibalism can provide a nutritional advantage despite substantial costs. In *Drosophila melanogaster* larvae, feeding is essential for successful metamorphosis, making food source decisions critical. We find that dead conspecifics act as attractive cues, but this attraction is weak for live larval groups. It is enhanced when dead larvae are injured or homogenized, or after repeated exposure. Larvae are also more attracted to dead individuals from other *Drosophila* or insect species. Testing live larvae individually revealed the strongest attraction, indicating that social context suppresses feeding-related attraction. Attraction to dead conspecifics depends on multiple chemosensory and mechanosensory modalities, highlighting the complex integration of social and nutritional cues in larval decision-making. Our results indicate that fly larvae flexibly balance attraction and avoidance based on social context to optimize resource use under ecological constraints.

**Short summary:** *Drosophila melanogaster* larvae flexibly balance attraction and social avoidance when encountering dead conspecifics. Attraction depends on carcass condition, prior experience, and social context, and is mediated by chemosensory, gustatory, and mechanosensory pathways. These findings reveal how larvae integrate multimodal social and nutritional cues to guide context-dependent feeding decisions.

## Introduction

Animals must flexibly adapt their behavior to survive in variable environments, responding to a wide range of cues, including signals from conspecifics. Social cues are often complex and less predictable than other environmental stimuli, and how the brain interprets and integrates them remains poorly understood. Under challenging conditions such as starvation, recognizing conspecifics becomes crucial to avoid maladaptive behaviors such as cannibalism. Although cannibalism can provide a critical nutritional advantage under resource scarcity, it carries substantial costs, including the potential spread of pathogens and reduced group survival.^1,2^

*Drosophila melanogaster* is a powerful model for dissecting the genetic, neural, and ecological foundations of social behavior.^3–5^ In adults, social behaviors such as mating and aggression have been extensively characterized, whereas in larvae, collective digging represents the best-described cooperative behavior.^6^ Under laboratory conditions, larvae develop in nutrient-rich environments that promote cooperative foraging. However, in nature, resources are patchy and unpredictable, and larvae may encounter non-optimal or even conspecific-based food sources. Previous studies have shown that larvae can resort to cannibalism when starved or raised on protein-poor diets, surviving on dead conspecifics or other insects.^7–11^ When given a choice, larvae preferentially consume distantly related species rather than their own, suggesting that cannibalism involves a decision-making process integrating species recognition and nutritional cues.^1,12^

We and others previously demonstrated that larval social behavior is highly context-dependent. ^6,13^ In the presence of food, larvae aggregate and cooperatively exploit protein-rich sources; in its absence, they disperse and avoid conspecifics. Here, we examine how larval groups balance attraction and avoidance when presented with dead, injured conspecifics as a food source. We show that attraction to dead larvae is generally weak but increases with prior exposure, physical damage to the carcass, or when dead individuals come from other *Drosophila* or insect species. Notably, individually tested live larvae are more attracted to dead conspecifics than larvae tested in groups. These results indicate that larvae can not only discriminate among species but also are aware of their social context and modulate feeding behavior accordingly.

We further identify the sensory basis of this behavior and show that attraction to dead conspecifics depends on chemosensory, gustatory, and mechanosensory inputs. These findings establish *Drosophila* larvae as a model for studying how multimodal social cues are integrated to guide complex, context-dependent decisions about conspecific interactions.

## Results

### Attraction to dead conspecifics

First, we asked if live larvae are contained by and attracted to dead conspecifics. Therefore, we exposed living *Drosophila melanogaster* larvae in groups of 15 to 10 dead conspecific larvae in a big petri dish arena (25cm x 25cm) filled with 2% agar as a crawling substrate for 15 minutes (Fig. 1A). The dead conspecifics were killed by short exposure to freezing temperatures (∼10 minutes). Afterwards, they were warmed up to room temperature and processed for the experiment, either left intact, stabbed with a needle, or torn apart with forceps (Fig. 1B). Groups of living fly larvae showed weak attraction to dead larvae. Intact dead larvae were the least attractive to live conspecifics, consistent with earlier studies (Fig. 1B). ^8,11^ Injured, dead victim larvae attracted more alive *Drosophila* larvae; however, only a few larvae engaged in cannibalistic feeding (see also ^9^). Dead, torn apart larvae increased attraction even more. The most attractive substance was normal fly food from food vials (Fig. 1B). Across all conditions, larvae chose to stay in the middle or to disperse within the first two minutes of the experiment (Fig. 1C). For further experiments, we decided to prepare dead larvae with stabbed injuries, which elicited an intermediate and reproducible attraction. To investigate whether larvae were actually feeding on dead conspecifics, we dyed the dead conspecifics with Brilliant Blue dye. Comparing the blue gut coloration of attracted and non-attracted, dispersing larvae revealed that most attracted larvae indeed ingest dead conspecifics, whereas dispersing larvae showed no blue coloration (Fig. 1D).

**Figure 1:**
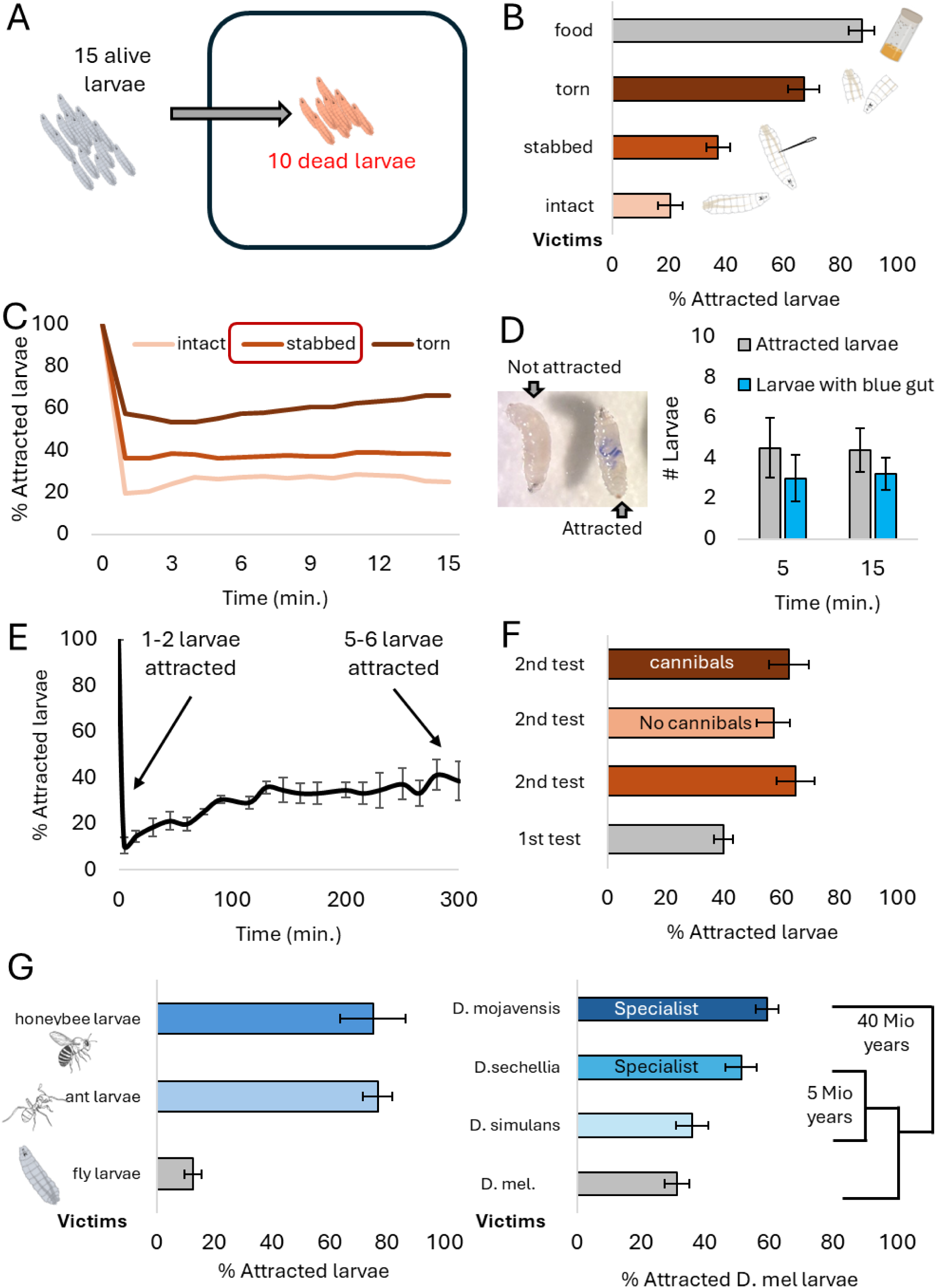
Fly larvae show weak attraction to dead conspecifics. (A) Experimental setup. An agar petri dish with 10 dead larvae in the middle of the arena. 15 alive larvae were added to the middle. After 15 minutes, live larvae in the middle are counted as attracted. (B) WT larvae’s preference for dead larvae processed in different ways (intact, stabbed, torn), and normal food. (C) Attraction over 15 minutes, same dataset as shown in (B). Attracted larvae were counted every minute. (D) Picture taken after an experiment with stabbed and dyed dead conspecifics (Brilliant Blue). Left: Non-attracted larva with no blue gut. Right: Attracted larva with blue gut. Attracted larvae and larvae with blue gut were counted after 5 and 15 minutes. (E) The experiment was set up as shown in (A) with live larvae recording for five hours. Larvae in the middle are counted every 15 minutes. (F) The same group of live larvae was repeatedly exposed to dead conspecifics (1st test/ 2nd test). Larvae in the middle (cannibals) and larvae that dispersed (No cannibals) were collected and retested independently. (G) Fly larvae were exposed to dead conspecifics or dead larvae from other insect species, leafcutter ant (*Atta vollenweideri*) or honeybee (*Apis mellifera*) larvae. *D. melanogaster* larvae were exposed to dead larvae from other *Drosophila* species (*D. mel. = Drosophila melanogaster*).

Next, we asked whether fly larvae might not be attracted to dead conspecifics because they represent an unfamiliar food source and thus cannot easily be identified as such. Therefore, we extended the experimental time and tested larval preference for five hours (Fig. 1E). Dead and alive larvae were placed in the middle of the arena, and the number of attracted larvae was counted every 15 minutes. In the first minutes of the experiment, larvae dispersed from the middle, and only a low percentage of larvae were attracted to the dead conspecifics, as described in previous experiments. During the five hours, only a few larvae returned to the dead conspecifics, with 40% of larvae being attracted at the end. Thus, not all live larvae get attracted over time to the unusual food source. We further asked whether repeated exposure to dead conspecifics can enhance attraction and tested larvae in two subsequent experiments (Fig. 1F). Between the experiments, larvae were housed on standard fly food. We found that when exposed to dead conspecifics for a second time, attraction to dead conspecifics was higher. We also collected attracted (cannibals) and non-attracted (no cannibals) larvae after the first exposure. In a second exposure experiment, both groups were tested independently; however, both showed higher and similar attraction, suggesting that the previous choice to leave or stay with the dead larvae did not affect a second choice (Fig. 1F).

To further test if fly larvae are generally not attracted to unknown food sources, we performed experiments with dead and injured larvae from different insect species and different *Drosophila* species (Fig. 1G). However, exposure to dead, injured larvae from leafcutter ants (*Atta vollenweideri*) or honeybee larvae (*Apis mellifera*) led to higher attraction than to fly larvae. This suggests that the unfamiliarity of the food is not the cause of the weak attraction, as the larvae had never been exposed to the larvae from other insect species. We further tested attraction to dead larvae from distinct *Drosophila* species (Fig. 1G). We tested two closely related sister species, the generalist *D. simulans* and the specialist *D. sechellia*, as well as a more distantly related specialist species, *D. mojavensis*.^14^ *D. melanogaster* live larvae showed weak attraction to their own species; however, attraction was higher to the two specialist species, *D. sechellia* and *D. mojavensis*. All species were grown on standard fly food; thus, we exclude the possibility that the specialist species were attractive due to differences in gut content. These results demonstrate that fly larvae can distinguish between different insect species and different *Drosophila* species, and that this information is relevant for choosing food sources.

### Dead conspecifics emit multisensory cues

To better understand the neural mechanisms underlying attraction to dead conspecifics, we asked how live larvae sense them. Therefore, we screened for the requirement of chemosensors using mutant lines and a genetic neural silencing method combining *UAS-kir* with sensory GAL4 driver lines (Fig. 2A-D). We previously controlled for normal locomotion in these genetically manipulated larvae and can thus exclude that any phenotypes are due to locomotion deficits.^13^ The ppk23 receptor is involved in attraction to larval deposits,^15^ egg avoidance,^16^ and avoidance of live conspecifics. ^13^ We discovered that the mutation of this receptor impairs attraction towards dead larvae (Fig. 2B, blue bars). We also tested for the requirement of olfactory receptors (ORs) and ionotropic receptors (IRs) using the anosmic mutant. This mutant carries mutations in the orco, IR8a, and IR25a genes (Fig. 2C, dark purple bar), however, it showed normal dead conspecific attraction. Notably, when testing these mutations individually, we found opposite phenotypes. Larvae without functioning olfactory receptors (orco mutant) showed reduced attraction to dead conspecifics, whereas larvae without functioning IR8a receptors showed enhanced attraction. This suggests that dead larvae might also emit chemical signals that induce avoidance. Silencing neurons expressing other IRs, such as IR25a or IR76b, which are broad IR co-receptors, did not affect dead conspecific attraction (Fig. 2C).

**Figure 2:**
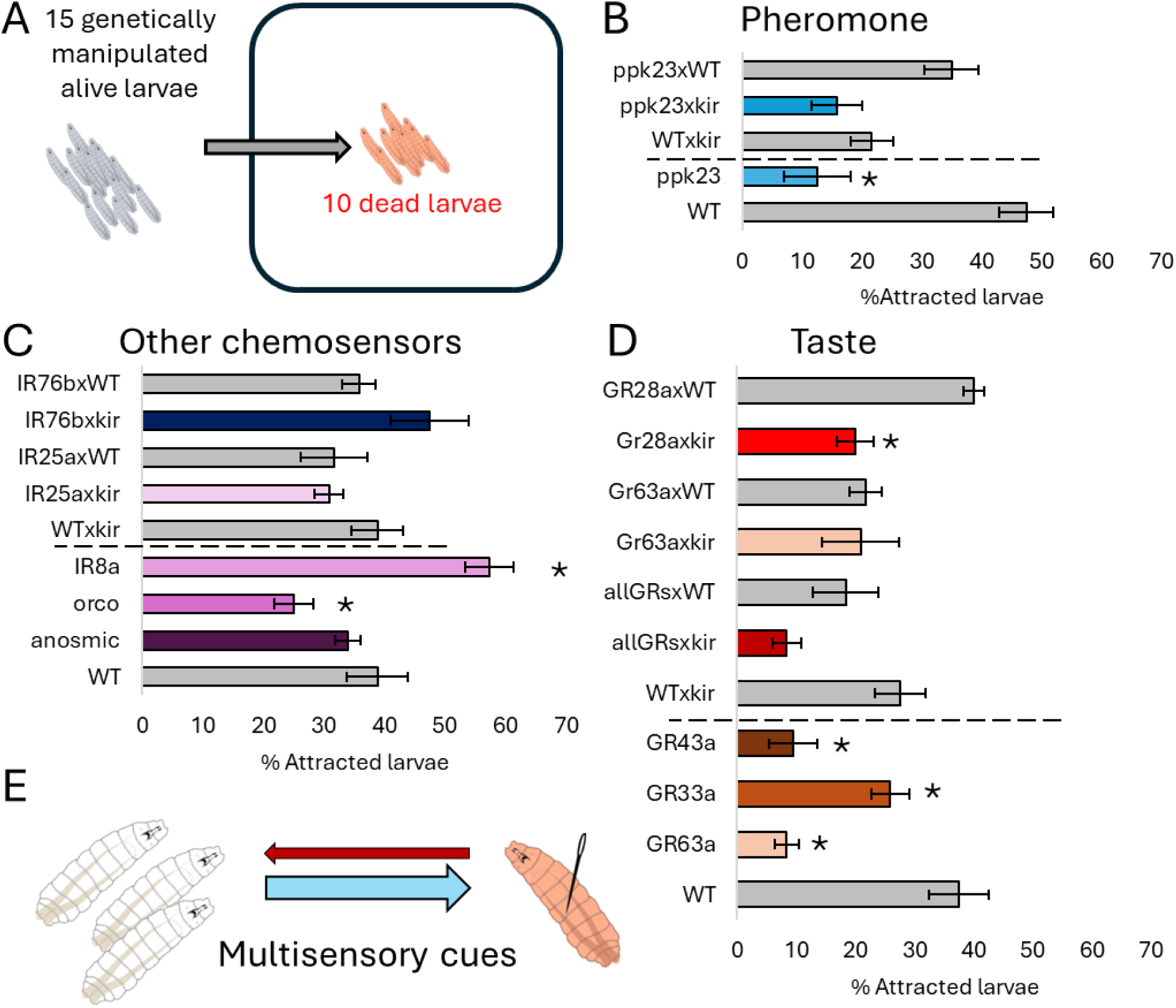
Attraction to dead larvae requires multiple chemosensors. 15 live larvae were exposed to 10 dead conspecifics for 5 minutes. The number of attracted larvae was counted at the end of the experiment. (A) Screen for the requirement of ppk23 receptor, IRs, and ORs. Larvae were counted after 5 minutes. (B) Screen for the requirement of GRs. (C) Dead conspecifics release mainly attractive cues, sensed via different modalities. Significance level was set to p<0.05.

As we observed that attracted larvae were feeding on dead conspecifics, we tested for the involvement of gustatory receptors (GRs, Fig. 2D). We found a tendency for reduced attraction when silencing many gustatory receptor neurons labelled by a broadly expressing allGRs-GAL4 line.^17^ Additionally, more specific mutations in GR63a, GR33a, and GR43a reduced dead conspecific attraction. Silencing GR28a-positive neurons with Kir also reduced attraction. Thus, the gustatory system is strongly involved in sensing and evaluating dead conspecifics. Overall, dead conspecifics emit multimodal chemosensory cues that are mainly attractive to live larvae (Fig. 2E).

### Social context modulates attraction to dead conspecifics

If dead conspecifics mainly emit attractive cues, why do alive larvae show only weak attraction and disperse in the arena? We previously showed that live larvae avoid other live conspecifics in the absence of food.^13^ Therefore, we hypothesized that the presence of other live larvae in the dead larva preference assay may also provide an aversive cue and tested individual live larvae for their dead conpsecific preference (Fig. 3A). Indeed, individual testing enhanced attraction to dead conspecifics from less than 20% to more than 60% (Fig. 3B). Individual larvae even showed enhanced attraction when we provided food-deprived dead conspecific victims (Fig. 3B). Thus, live larvae seemed to be interested in substances released by the dead larvae, other than food residuals in the gut. To test whether individual live larvae initially disperse and later return to dead conspecifics, we analyzed their attraction over time (Fig. 3C). Live larvae tested in groups of 15 showed avoidance in the first two minutes. Individually live larvae showed strong dead conspecific attraction over the whole experiment and did not initially disperse. Thus, the presence of live conspecifics decreases attraction to dead conspecifics within the first minutes of the experiment, and individual larvae, if attracted, stay with the dead conspecifics from the beginning.

**Figure 3:**
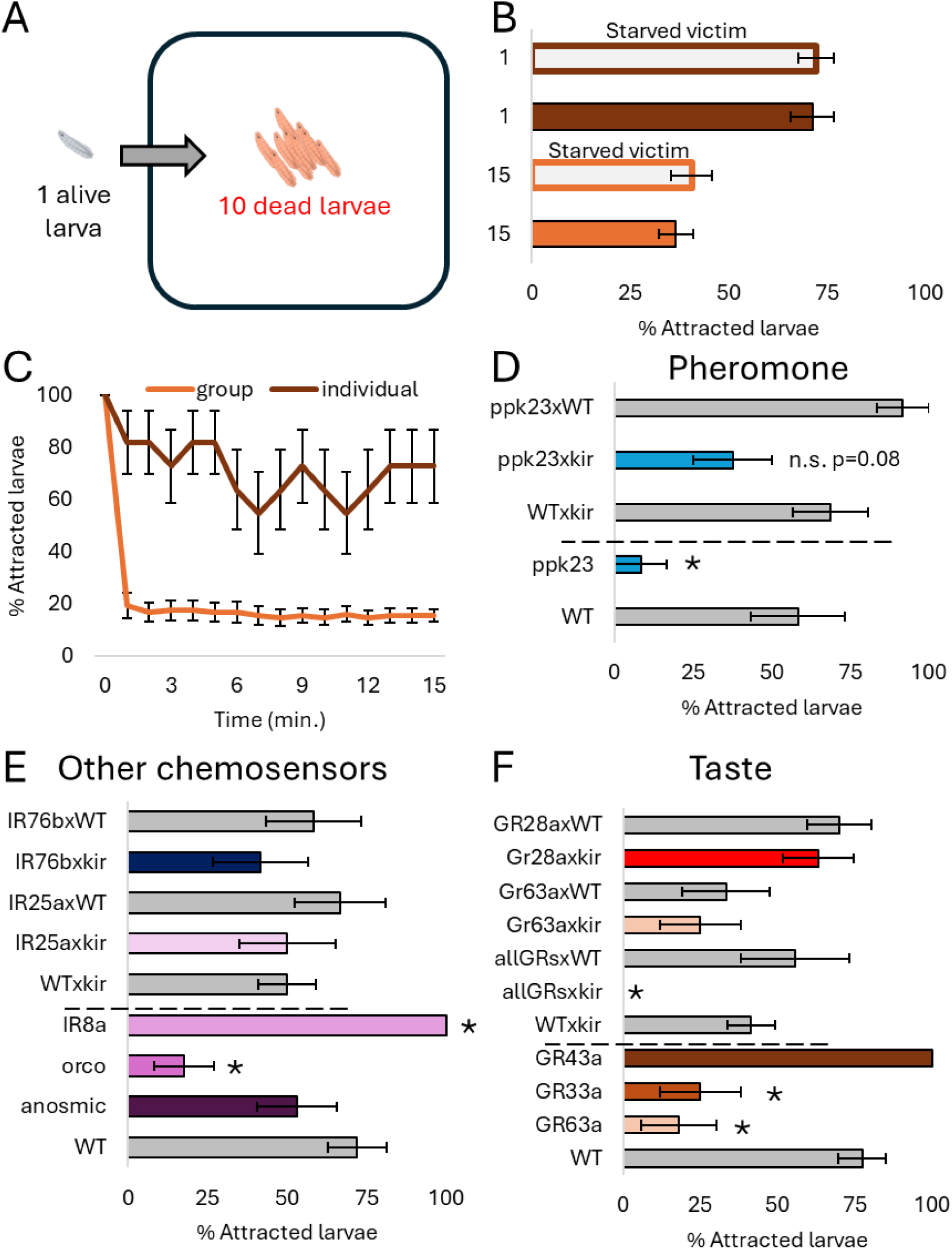
Individual live larvae show higher attraction to dead conspecifics. (A) Individual larvae were tested for their preference towards 10 dead conspecifics for 15 minutes. (B) Individual larvae have a higher probability of being retained by dead conspecifics. Single larvae and groups were exposed to fed or starved dead conspecifics. (C) Individual larvae stayed close to dead conspecifics over the time course of a 15-minute experiment. (D/E/F) Screen for the requirement of sensory neurons, including the pheromone ppk23 receptor (D), ionotropic and olfactory receptors (IRs/ORs) (E), and gustatory receptors (GRs) (F). Larvae were counted after 5 minutes.

We repeated the genetic screen for chemosensory requirement in the individual assay using the same mutant lines and genetic ablation methods (Fig. 3D-F). We confirmed several of the group-assay phenotypes. Testing individuals, the mutation of the ppk23 receptor also impairs attraction towards dead larvae (Fig. 3D, blue bars). We also found a strong tendency for reduced attraction when silencing ppk23-positive neurons. We could further verify that ORs and GRs are required for the attraction to dead conspecifics and that IR8a mutants showed enhanced attraction (Fig. 3E). For GRs, we could confirm that GR33a and GR63a mutants showed reduced attraction; however, silencing GR63a-positive neurons did not reveal a phenotype, similar to what we found in the group assay (Fig. 3F). GR43a mutant individual larvae showed the highest attraction (100%) to dead conspecifics, contrary to what we found in the group assay (compare to Fig. 2B). Silencing of GR28a neurons did not reveal a phenotype in the individual assay.

Here, we tested a single genetically modified larva for its attraction to dead conspecifics; thus, we specifically tested for sensory systems involved in sensing dead larval cues. The results confirmed that dead conspecifics emit multimodal sensory cues that are mainly attractive to live larvae (Fig. 2C).

### Mechanosensory cues are involved in dead conspecific sensing

In our dead conspecific preference assay, live fly larvae are placed in the middle of the dead conspecifics and touch them and each other before dispersing in the arena. Therefore, we tested the requirement for different mechanosensors using mutant lines and by genetically ablating sensory neurons, as previously described (Fig. 4). Most of the individual genetically modified live larvae were attracted to dead conspecifics, showing no impairment, likely because they encountered only dead conspecifics. Nanchung (NaN) mutants had reduced attraction. NaN is a mechanosensory channel required for vibration sensing in chordotonal neurons (Fig. 4A) ^18,19^ and soft food texture sensing in adults. ^20,21^

**Figure 4:**
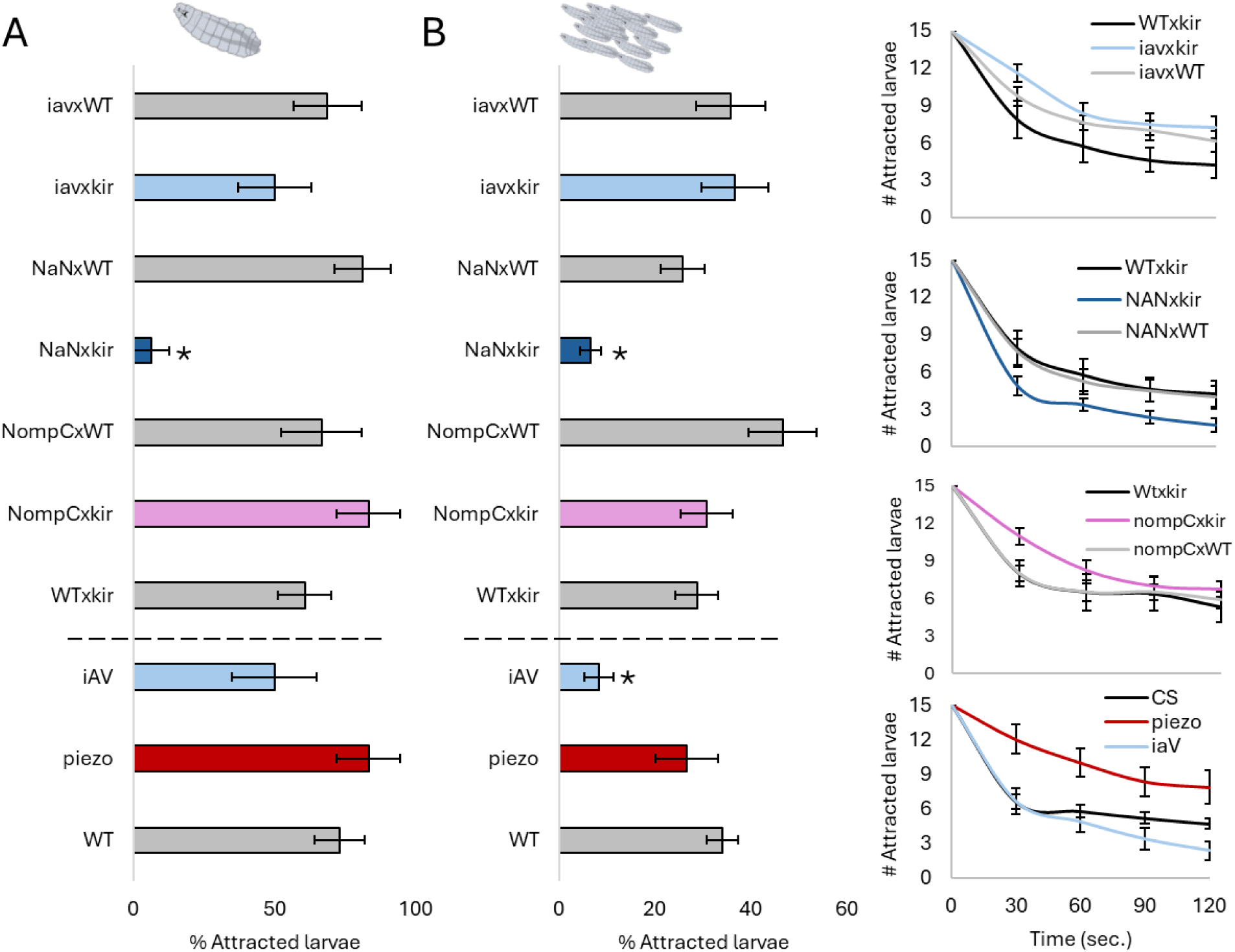
Dead and alive larvae provide mechanosensory cues required for attraction and avoidance. 15 live larvae or individual larvae were exposed to 10 dead conspecifics for 5 minutes. The number of attracted larvae was counted at the end of the experiment (individuals/ groups) or every 30 seconds (groups). (A) Screen for the requirement of mechanosensors in live, individually tested larvae. (B) Screen for the requirement of mechanosensors in live larval groups of 15. Attraction of larval groups over the first two minutes of the experiment reveals that piezo mutant larvae and NompC channel silencing delay dispersal of larval groups away from dead conspecifics.

Mechanosensory cues may be more prominent and relevant in the group assay, where live larvae interact with one another. Therefore, we also tested groups of genetically modified live larvae in the dead conspecifics assay (Fig. 4B). We consistently found that NaN mutants also disperse faster and show less attraction to the dead conspecifics. This is already noticeable in the first two minutes of the experiment. These results suggest that larvae sense dead larvae via mechanosensory cues, and this sensation is required for attraction. In the group assay, we additionally found reduced attraction with the inactive (iaV) mutant. Both channels, NaN and iaV, are often coexpressed and required for vibration sensing.^18,19^

Counting the number of attracted live larvae at the end of the experiment did not reveal further impairments. However, we also analyzed the preference over time for the group experiments (Fig. 4B, bar graphs) and found delayed dispersal in the piezo mutant and NompC (No mechanosensory potential channel) silenced larvae. A delayed avoidance response suggests that these channels are required for live conspecific avoidance. Neither manipulation showed a phenotype when tested in individuals, as most larvae remained in the center. We exclude that these phenotypes are due to locomotion defects, as we previously controlled for normal locomotion in these manipulated larvae by testing them in an assay without any stimulation.^13^ NompC is expressed in chordotonal organs and required for soft touch and vibration sensing in larvae.^19,22^ The mechanotransduction channel Piezo is involved in sensing harsher touch in larvae.^22,23^

## Discussion

Social behavior in *Drosophila melanogaster* larvae arises from competing drives of attraction and avoidance. We found that live larvae are attracted to dead conspecifics in a context-dependent manner. Attraction depends strongly on the preparation of the carcass—injured or homogenized larvae elicit stronger attraction than intact individuals—and is higher for dead larvae from other insect or *Drosophila* species. Larvae also adapt to repeated exposure: individuals that previously encountered dead conspecifics show increased attraction during subsequent encounters, although this preference is not individual-specific. The presence of live conspecifics induces an aversive response in larvae; therefore, they display the strongest attraction to dead conspecifics when tested alone. We further show that attraction to dead conspecifics relies on multimodal sensory input, requiring chemosensory, gustatory, and mechanosensory pathways, with distinct mechanosensory components contributing under group conditions. Together, these findings indicate that the decision to approach or avoid a dead conspecific involves complex integration of external cues, prior experience, and social context.

### Flexible exploitation of a non-optimal food source

Dead *Drosophila melanogaster* larvae act as attractive cues for conspecifics, likely reflecting their value as a potential food source when external resources are limited. This attraction is strongest in isolated larvae but diminishes in groups, suggesting that social context modulates the balance between attraction to food-related cues and avoidance of conspecifics. Our results indicate that this behavior is plastic and shaped by developmental experience. It has been shown that larvae reared under chronic malnutrition display enhanced cannibalism.^8^ Developmental crowding could suppress it, as repeated encounters with conspecifics might reduce the payoff of cannibalistic behavior.^24^ However, we find that larvae with previous exposure to dead conspecifics showed stronger attraction to carcasses than naïve individuals. We also previously found that rearing conditions affect social avoidance: isolated larvae avoid conspecifics more than group-housed larvae in non-nutritive contexts.^13^ Together, these findings suggest that larval social behavior is dynamically tuned by experience and environmental conditions, and that crowding might enhance cannibalistic feeding. Such behavioral plasticity may optimize resource use and minimize social conflict in fluctuating environments, balancing the benefits of conspecific exploitation with the costs of competition and kin harm.

We also found that larvae tested in groups approached dead heterospecifics or other insect species more often than dead conspecifics. This pattern implies that larvae can discriminate between conspecific and heterospecific cues and preferentially exploit non-conspecifics when food is scarce. Comparable kin discrimination has been reported in egg cannibalism, where larvae consume unrelated eggs more readily than kin eggs, likely to avoid indirect fitness costs.^12^ Species discrimination in insects often depends on cuticular hydrocarbons (CHCs)^25^, which differ among *Drosophila* species.^26^ The higher attractiveness of injured compared to intact larvae (Fig. 1B) further suggests that cuticular or hemolymph-derived compounds modulate attraction and avoidance. Notably, the strongest attraction to dead conspecifics occurred when larvae were tested individually, implying that social avoidance cues from live conspecifics compete with the attractive cues from dead individuals.

### Dead conspecific cues

Injured or homogenized *Drosophila* larvae elicit stronger attraction to live larvae than intact dead conspecifics (Fig. 1B). Exposure of hemolymph and internal tissues appears to provide potent attractive cues. Consistent with this, larvae are known to be attracted to conspecific deposits,^15^ a behavior mediated by the ppk23 receptor—one of the receptors we identified as necessary for attraction to dead conspecifics. This suggests that ppk23 contributes to species-specific recognition and mediates attraction in food-related contexts. In contrast, our previous work showed that ppk23 is required for social avoidance in the absence of food cues,^13^ indicating that its function is context-dependent, shifting between attraction and avoidance depending on the presence of nutritive signals.

Attraction to dead conspecifics also depends on multiple chemosensory receptors. Injured larvae release hemolymph and tissue-derived compounds, which likely activate many receptors. Mutating olfactory or specific gustatory receptors reduced attraction. GR33a mutants, which lack a receptor involved in bitter taste and male–male courtship inhibition,^27,28^ showed impaired attraction. GR33a has also been implicated in larval electrosensation.^29^ Similarly, mutants in GR63a — a receptor mediating CO₂ sensing in adults and larvae ^30^— displayed reduced attraction, implying that CO₂, electrical or chemical cues may together influence larval social and feeding behaviors.

Testing receptor function in individually tested larvae mostly confirmed the sensory pathways required for attraction to dead conspecifics. Chemosensory mutants largely recapitulated group-tested phenotypes, whereas mechanosensory phenotypes were reduced. Only silencing of neurons expressing the NaN receptor—required for gentle touch and vibration sensing ^19^ and for food texture detection in adults ^20^—significantly reduced attraction. Across experiments, neuronal silencing via the GAL4/UAS system generally produced weaker phenotypes than receptor loss-of-function mutations, likely reflecting incomplete GAL4 expression or limited coverage of all receptor neurons. These findings suggest that larvae evaluate not only chemical but also textural properties of dead larvae, using mechanosensory input to recognize dead conspecifics as potential food.

### Alive conspecific cues

Larval groups showed weak attraction to dead conspecifics and a tendency to disperse, suggesting that larvae must balance attraction to dead individuals with aversion to live conspecifics.^13^ The sensory requirements underlying this group-level behavior largely overlap with those identified in individually tested larvae, including ppk23, olfactory receptors (ORs), and gustatory receptors (GRs). However, silencing GR28a or mutating Gr43a produced phenotypes only in grouped assays. This may reflect the stronger influence of social avoidance in groups, which can reveal subtle sensory contributions that are redundant or undetectable in individual conditions. Gr43a mediates sweet perception,^31^ whereas GR28a is required for ribose and RNA sensing in larvae,^32^ suggesting that nutrient-related signals compete with social context to shape attraction.

In groups, living larvae encounter one another, potentially generating distinct mechanosensory input compared to interactions with dead larvae. Manipulating mechanosensory pathways revealed that silencing NaN-GAL4–expressing neurons or mutating *iav (inactive)* both reduced attraction to dead conspecifics. These two mechanosensory channels are largely coexpressed^19^ and appear necessary for detecting the physical cues associated with dead individuals. Temporal analyses further showed that *piezo* mutants and larvae with silenced NompC-expressing neurons remained longer near dead conspecifics and exhibited delayed dispersal. Both receptors are required for touch sensation, with Piezo mediating noxious mechanical touch and NompC responding to gentle mechanical stimuli.^22,23^ Impaired detection of live conspecifics in these mutants could thus reduce social avoidance, consistent with our previous findings that NompC-silenced larvae stay closer together in the social avoidance assay.^13^ Importantly, none of the mechanosensory manipulations affected general locomotion in empty arenas, indicating that these effects are specific to social and tactile processing. NompC may represent a conserved mechanosensory channel underlying social spacing, as it also contributes to social interaction in adult flies.^3^ These results highlight that the integration of chemosensory and mechanosensory inputs enables larvae to weigh competing cues of attraction and avoidance in social contexts.

Our findings suggest that dead and injured conspecifics are detected through a broader range of sensory modalities than live conspecifics.^13^ Group assay results uncovered the many sensory requirements for attraction to dead individuals, suggesting that when larvae face a conflict between avoiding live conspecifics and approaching dead ones, additional sensory input from dead larvae is required to overcome social aversion. How larvae integrate and represent such multimodal social information within the brain remains an open question. By identifying the key sensory modalities involved in recognizing live and dead conspecifics, our study provides a foundation for dissecting the neural pathways underlying social decision-making in *Drosophila* larvae.

More broadly, understanding how social stress shapes cannibalistic attraction and avoidance behaviors offers a window into how the nervous system evaluates competing social and survival-related cues. These findings may provide insight into conserved neural mechanisms by which organisms, including humans, regulate behavioral responses to social stress, anxiety, and aggression.

## Resource availability

### Lead contact

Request for further information and resources should be directed and will be fulfilled by the lead contact, Katrin Vogt.

### Materials availability

Fly strains are available from the *Drosophila* Bloomington Stock Center (see Table 1) or upon reasonable request from the corresponding author.

### Data and code availability

Raw data for all figures of the manuscripts is available in the attached Excel sheet (see Supplement).

## Acknowledgements

The authors would like to thank Armin Bahl and Einat Couzin-Fuchs for valuable discussions. We also thank the Thum lab, the Sprecher lab, and the Galizia lab for fly lines.

## Author contributions

Conceptualization, A.E., D.W., K.V.; methodology, investigation and analysis, A.E., D.W., N.T., R.P., and K.V.; writing, K.V.; supervision, K.V., and funding acquisition, K.V.

## Star Methods

### Animal stocks and husbandry

Flies were raised on a standard cornmeal diet in incubators at 25°C and 60% humidity on a 12-hour light/dark cycle. Adult flies were allowed to lay eggs for 48 hours and then removed from the vials. The eggs were allowed to develop for 4 to 6 days. Larvae of middle to late second instar stages were used for experimentation.

### Transgenic lines

Mutant fly lines and GAL4/UAS crosses for experiments were maintained as mentioned above and crossed with a specific number of animals to maintain similar crowding during development. For a single cross, at least 10 GAL4-driver line males and 25 UAS-kir2.1 female virgins were collected and allowed to mate. Control crosses for both the UAS-kir2.1 and the GAL4-driver line crossed to WT were also made with a similar sex ratio. Wildtype experiments were performed with the Canton-S strain. Stocks were obtained from the Bloomington *Drosophila* Stock Center or as a courtesy from other labs (BDSC, see Table S1).

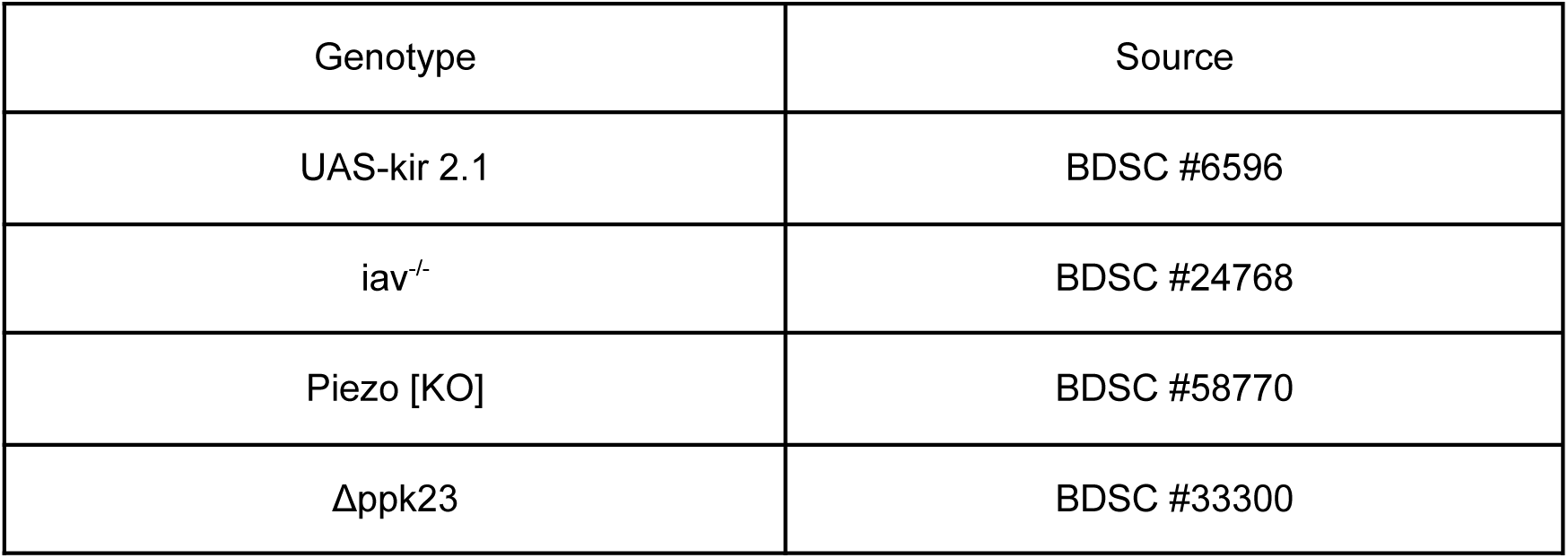
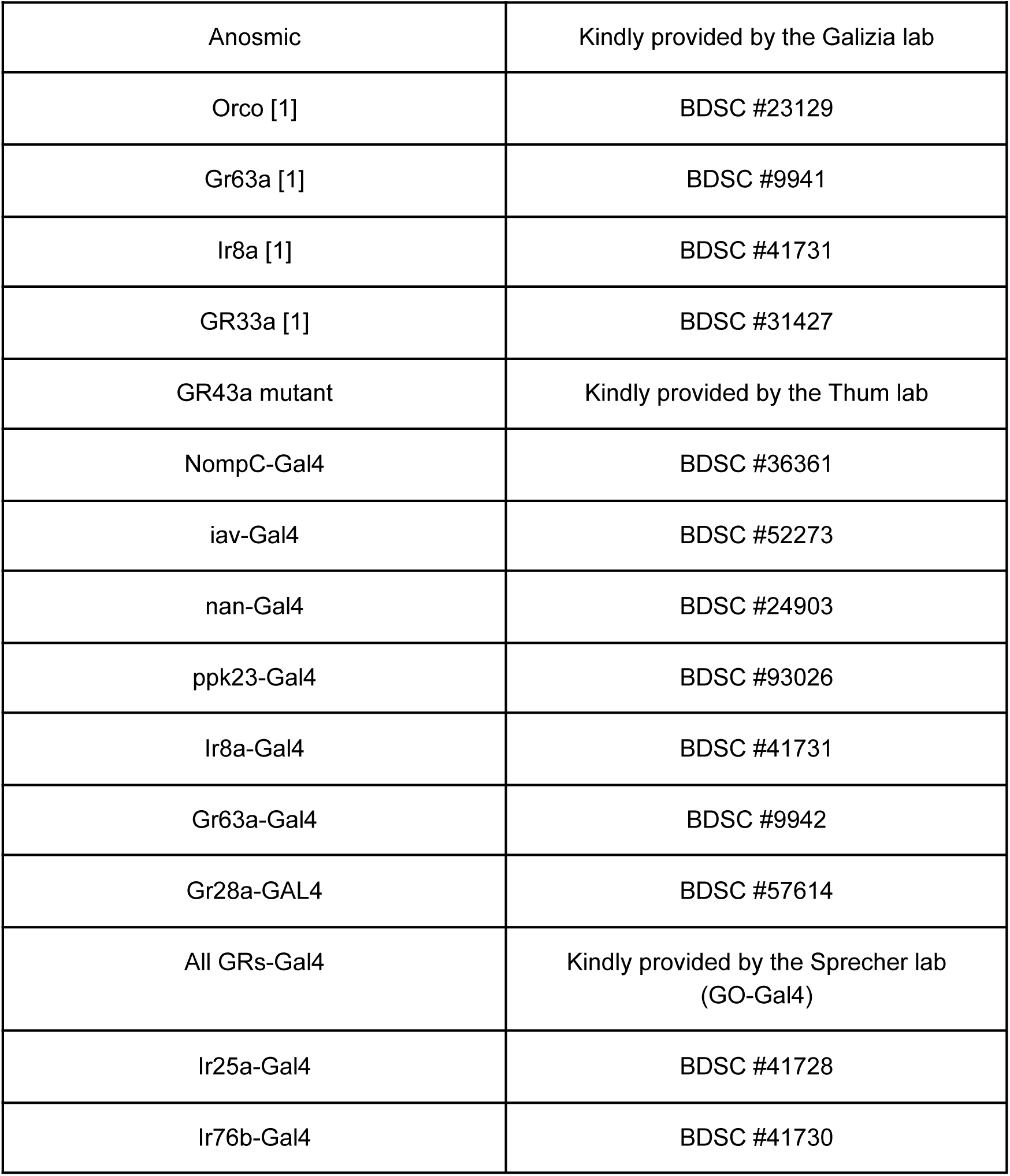

### Behavioral Experiments

#### Preparation of dead conspecifics

Larvae of instar stage three (L3) were removed from the fly vials and washed in distilled water to remove all traces of food. Clean larvae were placed in a small petri dish and kept in the freezer for at least 10 minutes. Dead and frozen L3 larvae were thawed and then used as dead conspecifics. They were tested either intact, several times poked with a sewing needle (standard procedure for most experiments), or torn apart with forceps. Prepared dead conspecifics were placed in the center of a 25 cm x 25 cm assay plate filled with 100 ml of a 2% agarose substrate. Experiments were performed immediately; one group of prepared dead conspecifics was used for up to three experiments, but not longer than 1 hour.

For the investigation of cannibalistic feeding, L3 larvae were first soaked in 3% Brilliant Blue water solution and then washed and placed in the freezer. Dead conspecifics with dye were handled as explained before and placed in the center of the agar plate. Live conspecifics were added, and at the end of the experiment, the number of attracted larvae was counted, and the number of larvae with a blue gut was counted.

#### Dead conspecific preference

Either a single live larva or a group of 15 live larvae was placed in the prepared pile of dead conspecifics and then allowed to roam freely on the 25 cm x 25 cm assay plate. The arenas were maintained at a room temperature (22°C) and placed in a light-tight box. Larval behavior was recorded for 5-15 minutes with a Basler camera (acA2040-90umNIR) and lens (Kowa Lens LM16HC F1.4 f15mm1”) at 1 fps with a red light filter (Edmund optics #89-837) above the arena. The arena was illuminated from below with red infrared LEDs (Solarox® LED strip infrared 850 nm).

For all behavioral experiments, middle to late second-stage instar live larvae (L2) of a similar size (4-6 days after egg laying) were used. Larvae were removed from the fly vials and washed in distilled water to remove all traces of food. To study individual behavior, a single larva was placed in the middle of the assay plate. To study group behavior, 15 larvae, unless stated otherwise, were placed in the center of the agarose plate. For repeated testing (Fig. 1D), larvae were collected after the experiment and stored in food vials to control for starvation effects. Before the second experiment, they were again washed in DI water and then placed in the arena with new dead conspecific larvae.

### Quantification and Statistical Analysis

The number of larvae retained in the pile of dead conspecifics was counted at the end of the experiment or at different time intervals during the experiment.

The percentage of larvae attracted to dead conspecifics was calculated as follows:

% Attracted larvae = (#larvae in dead conspecific pile / # all larvae)*100

To detect significant differences between experimental groups, we performed a non-parametric Mann-Whitney U test, which can be employed even with non-normally distributed data.

